# Cilia regulate meiotic recombination in zebrafish

**DOI:** 10.1101/2022.03.05.482955

**Authors:** Haibo Xie, Xiaosi Wang, Minjun Jin, Lanqin Li, Junwen Zhu, Yunsi Kang, Zhe Chen, Yonghua Sun, Chengtian Zhao

## Abstract

Meiosis is essential for evolution and genetic diversity in almost all sexual eukaryotic organisms. The mechanisms of meiotic recombination, such as synapsis, have been extensively investigated. However, it is still unclear whether signals from the cytoplasm or even outside of the cell can regulate the meiosis process. Cilia are microtubule-based structures that protrude from cell surface and function as signaling hubs to sense extracellular signals. Here, we reported an unexpected and critical role of cilia during meiotic recombination. During gametogenesis of zebrafish, cilia were specifically present in the prophase stages of both spermatocytes and primary oocytes. By developing a novel germ cell-specific CRISPR/Cas9 system, we demonstrated that germ cell-specific depletion of ciliary genes resulted in compromised double strand break repair, reduced crossover formation, and increased germ cell apoptosis. Our study reveals a previously undiscovered role for cilia during meiosis and suggests that extracellular signals may regulate meiotic recombination via this particular organelle.

## Introduction

Meiosis is probably the most key event in the life cycle of sexually reproducing eukaryotes. During the prophase of the first meiotic division, homologous chromosomes (homologs) of the maternal and paternal origins pair and synapse, which creates a context for the exchange of genetic information between homologs through an elaborate process, homologous recombination (HR). Meiotic recombination has been extensively studied in different model systems ranging from yeast, protists to mammals (Loidl, 2016; Zelkowski *et al*, 2019). In a classic HR model, DNA double-strand breaks (DSBs) are induced by the meiotic topoisomerase-like protein SPO11, then the phosphorylation of histone variant H2A.X is induced around the DSB sites. Meanwhile, DSB ends are processed and the resulting single strand overhangs are bound by recombinases including Dmc1 and Rad51. The broken DNA ends and these proteins form nucleofilaments and further invade the homologous template for homology searching and HR. Dmc1-mediated homology searching also promotes the formation of the synaptonemal complex (SC), an evolutionarily conserved tripartite protein structure formed between paired homologs. The SC is indispensable for multiple meiosis processes, including enhancing the interhomolog recombination, ensuring obligate crossover (CO) formation, and thus, safeguarding the faithful segmentation of chromosomes (see (Zickler & Kleckner, 2015)). Depending on the resolution of the double holiday junction, repairing of DSBs will be channeled to MutL homolog 1 (Mlh1)-dependent pro-CO pathway or non-CO pathways. The crossover recombination event leads to the reciprocal exchange of alleles between two homologues, thus ensures the genetic diversity of the progeny (Handel & Schimenti, 2010).

One of the key aspects of the meiotic recombination is the reorganization of the chromosome during prophase to ensure homologous pairing. The formation of the telomere bouquet is an evolutionarily conserved mechanisms for promoting homologous pairing (Loidl, 2016). Telomeres are attached to the nuclear envelope (NE) and clustered to one side of the nuclear through the linker of nucleoskeleton and cytoskeleton (LINC) complex, which bridges the movement of chromosome to the dynein motor moving on the perinuclear microtubules (Burke, 2018). Intriguingly, telomeres are usually clustered at the NE near the microtubule organizing center (MTOC), suggesting a potential role of the microtubule network in promoting the homologous pairing (Elkouby *et al*, 2016; Sato *et al*, 2009; Sawin, 2005). Although the mechanisms of homologous recombination has been extensively studied, it is still an enigma whether the meiotic recombination can be modulated by signals from the cytoplasm or extracellular environment.

## Results and discussion

Cilia are microtubule-based structures that are nucleated from the basal body, a highly conserved structure acting as the MTOC. The sperm flagellum is a special type of long motile cilia that is essential for sperm motility (Fig 1A). Unexpectedly, when examining the development of zebrafish sperm flagella, we observed a special cilia-like structure that were present in the spermatocytes (Fig 1A). Double immunostaining with anti-acetylated tubulin (a cilia marker) and SYCP3 (a lateral element of SC) antibodies showed that these ciliary structures were initially formed at leptotene stage, and gradually elongated until diplotene stage (Fig 1A). The diplotene cilia was about half length of sperm flagella (Fig 1A-B). Interestingly, these cilia were disappeared at diakinesis stage, implying that they are not the rudiments of sperm flagella. Further immunostaining with the centrosomal marker, anti-γ tubulin antibody, indicated that these microtubules were nucleated from one of the centriole (Fig 1C, S1). Moreover, these cilia can also be labeled with anti-glutamylated tubulin antibody (Fig S2A). Whereas, although sperm flagella were positive for both mono-or poly-glycylated tubulin antibody, these spermatocyte cilia were lack of this kind of tubulin modifications (Fig S2B). Furthermore, ultrastructural analysis showed that these cilia had a “9+0” axonemal configuration, in contrast to the “9+2” arrangement of sperm flagella (Fig 1D-F’). Finally, this special type of cilia was also present in the leptotene and zygotene stages of primary oocytes (Fig S3).

**Fig 1.**
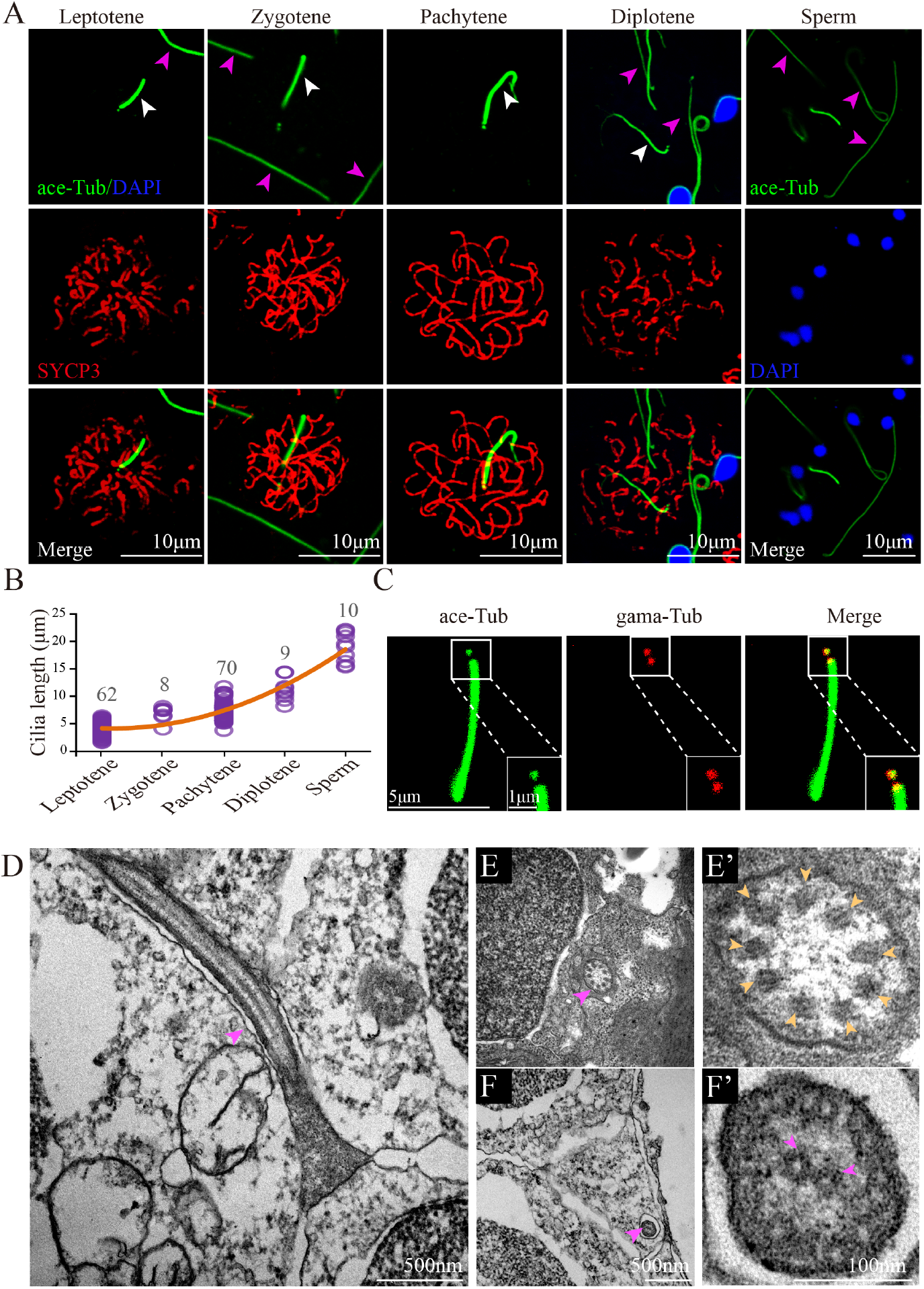
Cilia of primary spermatocytes in meiosis. (A) Confocal images showing cilia (white arrowhead) and sperm flagella (pink arrowhead) labeled with anti-acetylated tubulin (ace-Tub, green) antibody. The synaptonemal complexes were labeled with SYCP3 (red) and nuclei were counter-stained with DAPI in blue. (B) Statistical results showing the length of cilia in different stages of primary spermatocytes and sperms. (C) Confocal images showing cilia and basal bodies labeled with anti-acetylated tubulin (ace-Tub, green) and anti-γ tubulin (gama-Tub, red) antibodies on primary spermatocyte. (D-F’) TEM results showing the ultrastructure of primary spermatocyte cilia (D-E’) and sperm flagella (F-F’). Cross section showing the “9+0” configuration of spermatocyte cilia (E’).

The presence of these cilia during prophase I of the first meiotic division implies that cilia may participate in the regulation of homologous recombination. As cilia are essential for early zebrafish embryos development (Song *et al*, 2016), we seek to design a strategy to knockout these cilia specially in the spermatocytes. To reach this goal, we developed a germ cell-specific CRISPR/Cas9 system by generating a primordial germ cell (PGC)-specific Cas9-transgenic line, *Tg(kop:cas9-p2a-egfp-UTRnanos3)* (for abbreviation: *Tg(kop:cas9-UTRnanos3)*) (Fig 2A), in which a Cas9-2A-EGFP expression cassette was placed under the control of the PGC-specific *askopos* (*kop*) promoter and *nanos3* 3’-UTR (Xiong *et al*, 2013). Since the Cas9 protein was expressed only in the PGCs of *Tg(kop:cas9-UTRnanos3)* embryos, injection of gRNA into the embryos would not result in somatic mutation but in PGCs-specific gene knockout. Further crossbreeding of the injected F0 fish could generate maternal zygotic mutants in the F1 generation, even if the zygotic mutants were lethal (Fig S4A). To test the efficacy and penetrance of PGCs-specifically expressed Cas9 in the *Tg(kop:cas9-UTRnanos3)* embryos, we first injected a validated sgRNA targeting *tcf7l1a* (Zhang *et al*, 2020), whose maternal zygotic mutants (MZ*tcf7l1a*), but not maternal (M*tcf7l1a*) or zygotic mutants (Z*tcf7l1a*), shows a headless phenotype (Kim *et al*, 2000). Crossing two high-efficient germline mutated females and males resulted in a typical headless phenotype in more than 50% of the F1 embryos (Fig S4B, C), suggesting the high-efficiency of PGCs-expressed Cas9. Similarly, we were also able to generate MZ*pou5f3* mutants which exhibited severe developmental defects (Reim & Brand, 2006; Schulz & Harrison, 2019) (Fig S4 D,E). These data demonstrate that the *Tg(kop:cas9-UTRnanos3)* line expresses a highly efficient Cas9 protein which enables germ cell-specific gene knockout. Noticeably, albeit with high efficiency, the penetrance of mutations in the gametes was not 100%, which may be due to the temporal presence of sgRNAs.

**Fig 2.**
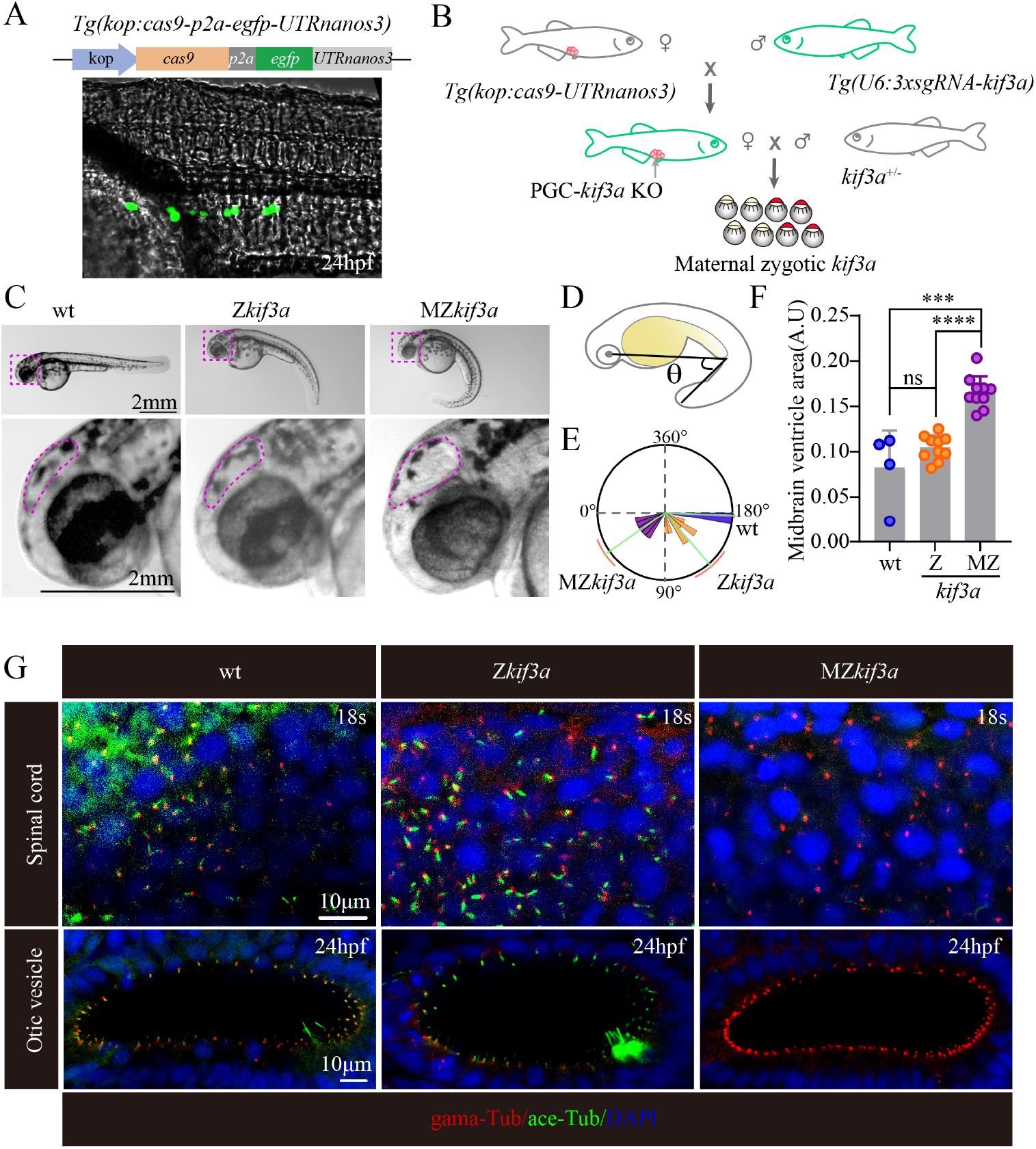
Generation of MZ*kif3a* mutants via germ cell-specific expression of Cas9. (A) Bright field image showing a 24 hpf *Tg(kop: cas9-p2a-egfp-UTRnanos3)* zebrafish embryo with EGFP fluorescence in the PGCs. Top: Diagram of the construct for making this transgene. (B) Schematic diagram showing the strategy for generating germ cell-specific knockout mutants of *kif3a* (gcKO*kif3a*). (C) External phenotypes of 24 hpf Z*kif3a* and MZ*kif3a* mutants. The enlarged boxes were shown in the bottom and dots outline the midbrain ventricles. (D-E) Statistical analysis showing the increased body curvature severity as demonstrated by the reduced angles of MZ*kif3a* mutants. (F) Bar graph showing relative size of midbrain ventricles in different mutants as indicated. A.U. arbitrary unit. (G) Confocal images showing cilia in the spinal cord and otic vesicle of wild type, Z*kif3a* and MZ*kif3a* mutants as indicated. Cilia were labeled with acetylated tubulin in green and the basal bodies were stained with γ-tubulin in red. Nuclei were counterstained with DAPI.

To further improve the efficiency of gene editing in the PGCs, we generated a stable *Tg(U6:3xsgRNA-kif3a)* transgenic line, in which three sgRNAs targeting zebrafish *kif3a* was each driven by an individual U6 promoters in tandem. Kif3a is a plus-end motor protein crucial for cilia assembly (Zhao *et al*, 2012). We further crossed the *Tg(kop:cas9-UTRnanos3)* line with the *Tg(U6:3xsgRNA-kif3a)* line (Fig 2B) to generate the *Tg(kop:cas9-UTRnanos3; U6:3xsgRNA-kif3a)* double transgenic line. By crossing the double transgenic female with *kif3a* heterozygotic male, we have successfully generated maternal-zygotic *kif3a* mutants (MZ*kif3a*). Around 50% of embryos (340 of 707) displayed body curvature defects, strongly implying that Kif3a was deficient in almost all of the eggs produced from the double transgenics. Compared with Z*kif3a* mutants, MZ*kif3a* mutants displayed strong body curvature defects, together with the increased size of the brain ventricles, a typical feature of hydrocephalous (Fig 2C-F) (Feng *et al*, 2017). Further whole mount staining results confirmed that cilia were completely absent in the MZ*kif3a* mutants, while cilia were still formed in the spinal cord and otic vesicle of Z*kif3a* mutants at early stages due to the maternal effects of Kif3a (Fig 2G). As for the double transgenic males, although they developed to adulthood normally as the wildtype males, they could not fertilize wildtype eggs due to the germ-cell disruption of Kif3a (Fig 3A-C). The spermatozoa were able to be produced in the testis, while all the flagella were significantly shorter than those of control siblings (Fig 3D-E, S5A). These data also suggest that sperm flagellum can still be partially assembled in the absence of Kif3a.

**Fig 3.**
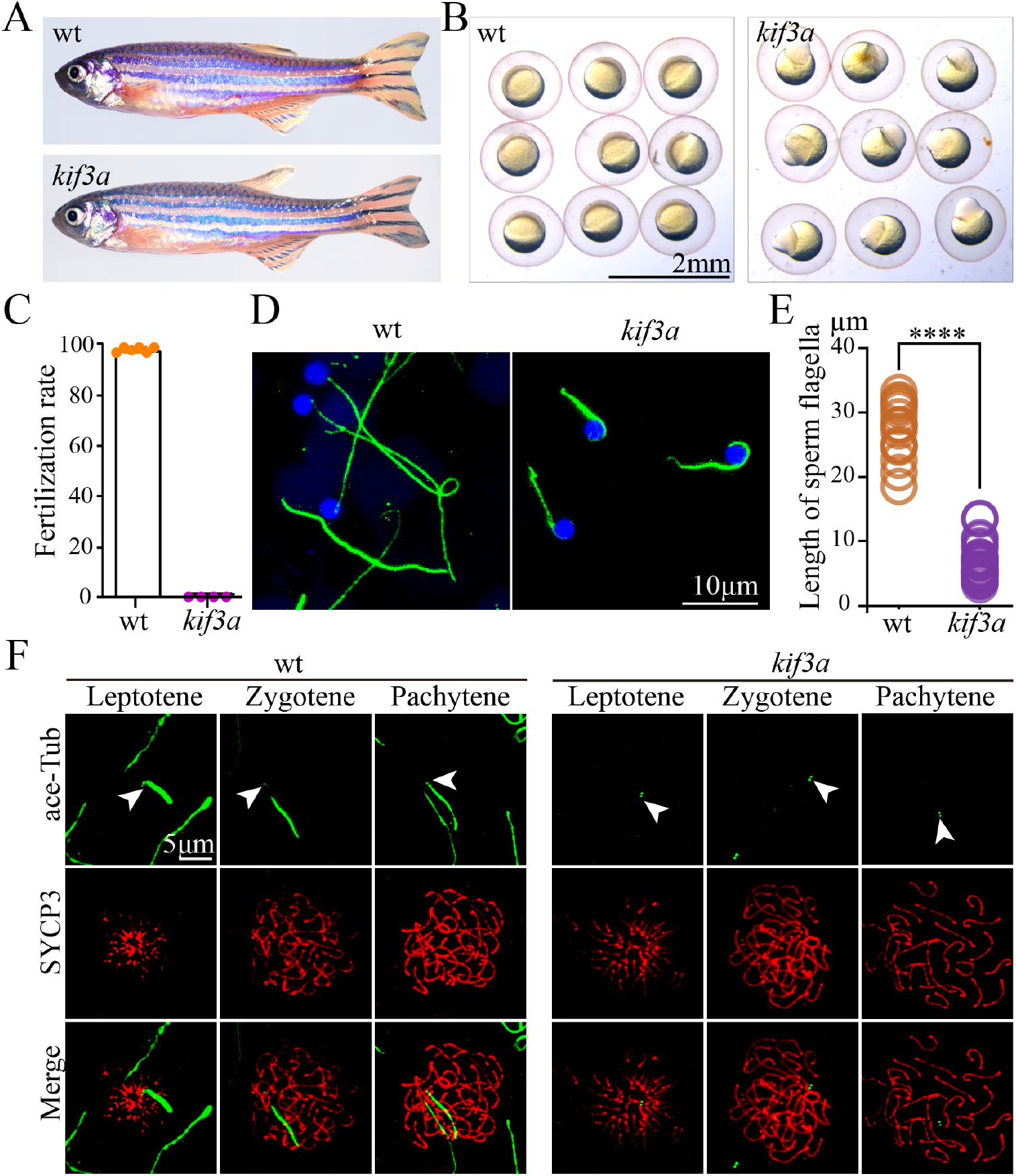
Phenotypes of germ cell *kif3a* conditional knockout mutants. (A) External phenotypes of wild type and *Tg(kop: cas9-UTRnanos3; U6:3xsgRNA-kif3a)* double transgenic male fish. (B) The embryos produced from wild type female crossed with control or double transgenic male at 6 hours after fertilization. (C) Dot plot showing the percentages of fertilization rates of wild type and double transgenic males as indicated. (D) Confocal images showing sperm flagella in wild type and double transgenic fish labeled with anti-acetylated tubulin antibody. (E) Statistical analysis of the length of sperm flagella in wild type and double transgenic fish. (F) Confocal images showing cilia in primary spermatocytes of wild type and double transgenic fish. Spermatocyte cilia (green) were absent in the double transgenic fish. The stages of spermatocytes were distinguished by the staining of SYCP3 (Red).

Since Cas9 protein was expressed in PGCs, it is rationale that *kif3a* gene should be disrupted at earlier stages. Indeed, cilia failed to form in the primary spermatocytes, and the staining of acetylated tubulin was only detected in the centrosomes (Fig 3F). Further staining with SYCP3 on the sections of testis showed that cilia were completely absent in all the earlier stage primary spermatocytes (Fig S5B), which further confirmed the nearly 100% efficiency in inducing germ cell-specific knockout of *kif3a* (gcKO*kif3a*) by the double transgenic approach (Fig 2B). Noticeably, strong signals were still present in those regions that were rich for spermatozoa, which were related to the remaining shorter sperm flagella (Fig S5B). Moreover, cilia in the primary oocytes were also absent in the double transgenic females (Fig S3).

During meiotic recombination, the formation of DSB provokes a rapid DNA damage response, leading to the phosphorylation of a histone H2A isoform at sites of DSBs. By using the anti-γ-H2A.X antibody, we can distinguish spermatocytes at both leptotene and zygotene stages. The staining of γ-H2A.X was gradually disappeared from mid-zygotene stage (Xie *et al*, 2020). The staining signals were similar between gcKO*kif3a* spermatocytes and control siblings at leptotene stage, while unexpectedly, the spermatocytes had strong γ-H2A.X staining throughout their nuclei at zygotene stage in gcKO*kif3a* mutants (Fig 4A-B). By counting the number of γ-H2A.X positive germ cells, we further found that the number of spermatocytes within each spermatocyst was comparable between the gcKO*kif3a* and control fish (Fig 4C), suggesting that loss of spermatocyte cilia did not affect the mitotic division of spermatogonium.

**Fig 4.**
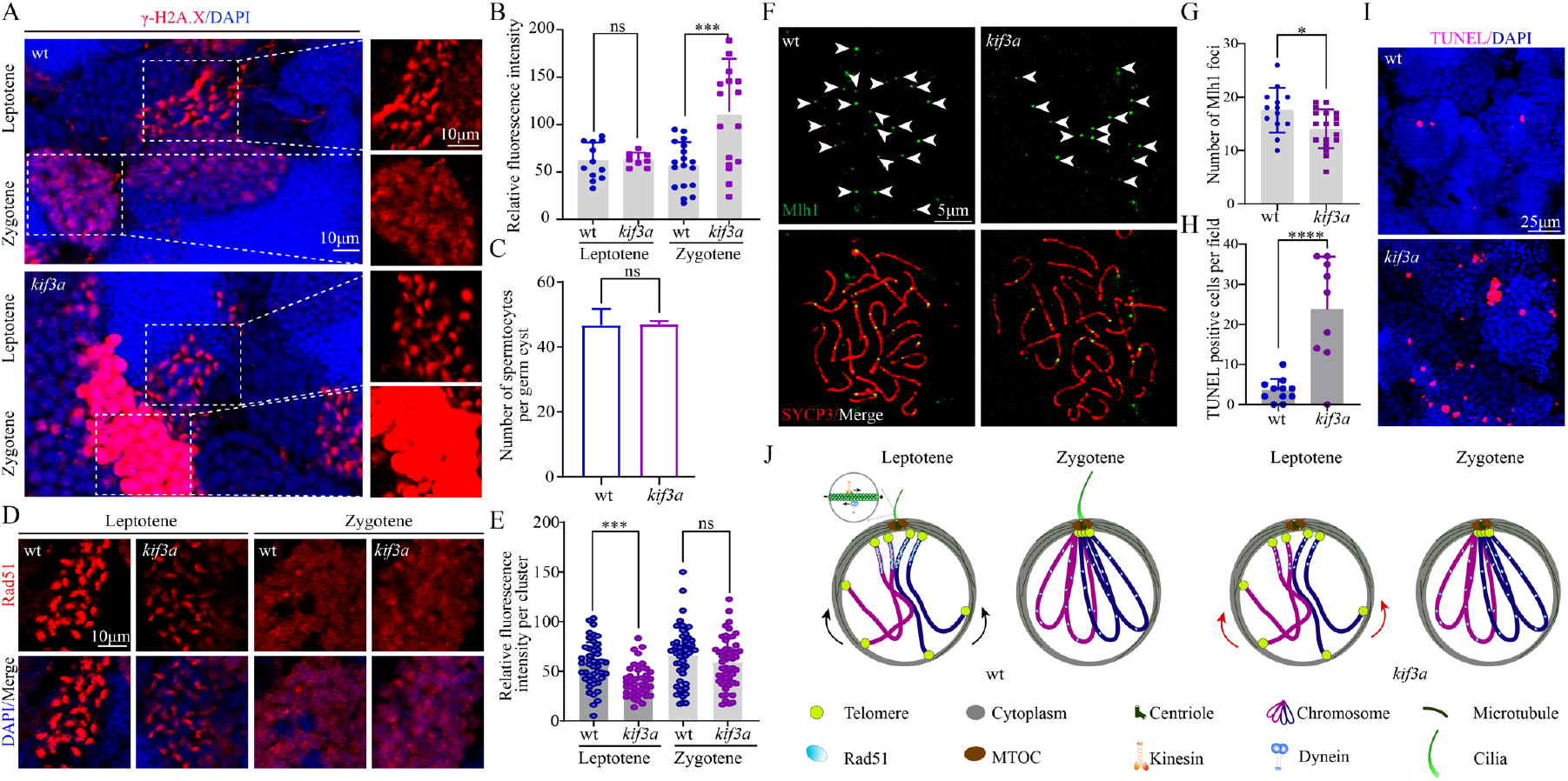
The repair of DSBs was compromised in the absence of spermatocyte cilia. (A) Confocal images showing the staining of γ-H2A.X (red) in the testis of wild-type and *Tg(kop: cas9-UTRnanos3;U6:3xsgRNA-kif3a)* double transgenic fish. (B) Statistical analysis showing relative fluorescence intensity of γ-H2A.X staining on leptotene and zygotene spermatocytes of wild type and double transgenic fish. (C) Statistical analysis of the number of spermatocytes per germ cyst in wild type and double transgenic fish. (D) Staining of Rad51 in the testis of wild-type and double transgenic fish. (E) Statistical analysis of relative fluorescence intensity of Rad51staining on leptotene and zygotene spermatocytes. (F) Staining of Mlh1 in primary spermatocytes of wild-type and double transgenic fish. (G) Statistical analysis of the Mlh1 foci number per spermatocyte. (H-I) Apoptotic cells stained by TUNEL assay in wild type and double transgenic fish. (J) Model illustrating the roles of spermatocyte cilia in DSBs repair. In the absence of cilia, less Rad51 proteins are recruited to the DSBs near the MTOC, which compromises the efficiency of DSBs repair.

The strong signals of γ-H2A.X staining suggest that DSB repair may be affected in the absence of spermatocyte cilia. Next, we checked the distribution of Rad51, one of the key recombinases recruited to the break sites for DSB repair. Compared with those of control cells, Rad51 signals decreased substantially in leptotene-stage spermatocytes of gcKO*kif3a* males (Fig 4D-E). Rad51 signals became weaker at zygotene stage and there were no difference in the staining of Rad51 between mutant and control siblings at later zygotene stage (Fig 4D). Rad51 is required for the formation of crossovers. Next, we evaluated the crossover events by investigating the number of Mlh1 foci on surface spread spermatocytes. The number of Mlh1 foci decreased significantly in the mutant spermatocytes (Fig 4F-G). Finally, TUNEL assay results indicated that mutant testis contained high number of apoptotic spermatocytes than those of control siblings (Fig 4H-I). Altogether, these data suggested that loss of spermatocyte cilia affected the DSB repair process, leading to decreased crossover formation and cell apoptosis.

Finally, to exclude the non-ciliary role of Kif3a during meiotic progression, we further generated *Tg(hsp70l:ift88-egfp)* transgenic line in which *ift88*, another essential gene for ciliogenesis (Tsujikawa & Malicki, 2004), was driven by a heat shock promoter. By performing routine heat shock, we were able to rescue *ovl*(*ift88)* mutants with this transgene (Fig S6A). Interestingly, the adult mutants can still survive for a short period even without heat inducement. By comparing mutants between heat induced and untreated fish, we found that the spermatocyte cilia were lost in the testis of non-heat-shocked controls (Fig S6B-C). Moreover, mutants generated from these non-heat-shocked adult females and heterozygotic males showed stronger body curvature and ciliogenesis defects, resembling those of MZ*ovl* embryos (Fig S7) (Huang & Schier, 2009). Similar to those of *kif3a* mutants, loss of *ift88* also caused abnormal cell apoptosis of the spermatocytes (Fig S6D-E), further confirming the role of cilia during meiosis processing.

In summary, here we reported a novel type of cilia existing in the primary spermatocytes and oocytes during meiotic recombination. By developing a germ cell-specific gene knock out approach, our study demonstrates that depletion of these early germ cell-specific cilia results in compromised processing of DSB repair, impaired crossover formation, and increased cell apoptosis as well (Fig 4J). It is noteworthy that oocytes and spermatozoa could be still produced in the absence of germ cell-specific cilia, albeit with cell apoptosis and flagellar defects in the spermatozoa. It is likely that this cilium may regulate DSB repair to ensure crossover formation, whereas it is not essential for the processing of meiosis. Cilia act as a cellular antenna to sense extracellular signals (Ishikawa & Marshall, 2011). Although we do not know yet the detailed mechanisms of these cilia in the regulation of meiotic recombination, our data provide an important message that meiotic recombination can be regulated by signals outside the nucleus. These cilia may function as signal hubs to sense signals outside the cell that help recruit DSB repair components, such as Rad51 and DMC1, to the DSB sites and ensure the HR processing. On the other hand, these cilia are nucleated from the basal body, from which the bouquet microtubule originates. It is possible that the cilia may supply forces for dynein/kinesin motors to support telomere movement and thus homolog pairing (Fig 4J). Further experiments are needed to dissect the precise signaling molecules underlying this intriguing organelle, and the germ cell-specific CRISPR/Cas9 system developed in present study may provide a powerful tool to investigate this process.

## Materials and methods

### Ethics statement

All zebrafish study was conducted according standard animal guidelines and approved by the Animal Care Committee of Ocean University of China and the Animal Care Committee of the Institute of Hydrobiology, Chinese Academy of Sciences.

### Zebrafish strains

All zebrafish strains were maintained at 14-hour light /10-hour dark cycle at 28.5°C. The plasmid for generating *Tg(kop:cas9-p2a-egfp-UTRnanos3)* transgenic fish was generated by replacing the *egfp* sequence in the pTol2(kop:egfp-UTRnanos3) construct (Xiong *et al*., 2013), with an in frame sequence, cas9-p2A-egfp, which encodes zebrafish codon optimized Cas9, p2A peptide, and EGFP. The plasmids for *Tg(U6x:3xsgRNA-kif3a)* transgene was generated by Golden Gate cloning as previously described (Yin *et al*, 2016). Three single guide RNAs (sgRNAs) targeting *kif3a* (sgRNA1, GGGAAAACGTTCACTATGGA; sgRNA2, GGCCAAACTCGACATGGAGG; sgRNA3, GAGGAGGTGAGAGATCTGTT) were each driven by distinct zebrafish U6 promoters. The *Tg(U6x:3xsgRNA-kif3a)* transgenic fish were further crossed to *Tg(kop:cas9-p2a-egfp-UTRnanos3)* transgenic line to get the double transgenic fish. The pTol2-*hsp70l:ift88-egfp* plasmid was constructed through Multisite Gateway cloning and injected into the homozygous *ovl* (*ift88*) mutants at one cell stage. The injected embryos were heat induced for three hours every day until adulthood for further analysis.

### Microinjection of single guide RNA (sgRNA)

For microinjection based primordial germ cell targeted gene knock experiment, single guide RNAs against *tcf7l1a* and *pou5f3* were GGAGGAGGAGGTGATGACCT and GGGTGAACTACTACACGCCA respectively, as previously described (Zhang *et al*., 2020). The sgRNAs were injected with a dosage of 80 pg per embryo into the *Tg(kop:cas9-p2a-egfp-UTRnanos3)* transgenic embryo at 1-cell stage.

### Chromosome spreading and immunostaining

Chromosome spreading steps were similar to previously published (Xie *et al*., 2020). Briefly, the dissected testis was first incubated in 50 µl 1 X PBS for 5 min, then transferred into 300 µl hypotonic solution (100 µl 1xPBS plus 200 µl double distilled water) for 25 min on an adhesive slide. After fixation with 4% paraformaldehyde (PFA) for 5 min, the cells were washed three times with PBS, blocked with blocking solution (10% goat serum,3%BSA,0.05% TritonX-100) for 30 min, then followed by regular antibody staining.

### Immunofluorescence and TUNEL assay

Zebrafish testes were first dissected and fixed in 4% PFA overnight at 4°C. After three times wash with PBS for 10 min each, the testes were infiltrated in 30% sucrose overnight at 4°C, then embedded with tissue freezing medium (OCT). Cryosectioning was performed using LEICA cryostat (CM1860). TUNEL assay was performed using the in situ cell death detection kit (Roche) according to standard protocols from the manufacturer. Images were collected with a Leica Sp8 confocal microscope.

To visualize cilia in the primary ovaries, zebrafish females about 1.5 month old were first euthanized with ice-cold water, and ovaries were dissected and transferred into 4% PFA overnight at 4°C. After three times wash with PBST (1 X PBS + 0.1% tween20), the ovaries were further blocked with Blocking Solution (0.3% TritonX-100, 10% Fetal Bovine Serum, 1xPBS) for 1.5h at room temperature. Antibody staining steps were similar to those of testes.

For immunofluorescence, the following antibodies were used: mouse anti-acetylated tubulin, rabbit anti-SYCP3, rabbit anti-γ-H2A.X, rabbit anti-RAD51, mouse anti-MLH1, mouse anti-γ-tubulin, rabbit anti-acetylated tubulin, mouse anti-glutaminated tubulin, mouse anti-monoglycylated tubulin and mouse anti-polyglycylated tubulin.

### Immunohistochemical staining

Zebrafish testes were dissected and fixed in 4% PFA overnight at 4°C. After gradual dehydration through 30%, 50%, 75%, 95% and 100% ethanol, the testes were embedded in JB4 embedding medium (Polysciences Inc., Warrington, PA, USA). Transverse sections through the testes were collected using Leica RM2235 microtome and stained with hematoxylin and eosin. Images were taken using a Leica DM2500 microscope.

### Ultrastructural analysis

Zebrafish testis were dissected and fixed overnight in 2.5% glutaraldehyde. The testis was washed three times in PBS for 10 min each, fixed again in 1% osmium tetroxide. After that, the testis was washed, gradually dehydrated to 100% acetone, and then embedded in Epon812 resin. Ultrathin sections were collected and stained with uranyl acetate and lead citrate. The sections were observed on a transmission electron microscope (JEM-1200EX) and imaged with Olympus Soft Imaging Solutions.

### Statistical analysis

All the confocal images were captured using Leica Sp8 confocal microscope. To compare the fluorescence signals between control and mutants, all the images were collected at the same parameters within each experiments. All experiments were repeated at least three times. Statistical analysis was performed using ImageJ, Microsoft Excel or GraphPad Prism 6 software. All data were presented as mean ±S.D. as indicated in the figure legends. A value of p<0.05 was considered statistically significant.

## Acknowledgement

We thank Dr. Feng Xiong at the China Zebrafish Resource Center for generating the pTol2(kop: cas9-p2a-egfp-UTRnanos3) construct. We thank Dr. Liangran Zhang, Wei Li and Ming Shao for providing DNA constructs and reagents. We also thank Dr. Miao Tian and Tao Qiu for their help during the preparation of this manuscript. This work was supported by the National Natural Science Foundation of China (Nos. 31991194, 32025037, 32125015), grant from the Fundamental Research Funds for Central Universities (NO. 201941004) and grant from the State Key Laboratory of Freshwater Ecology and Biotechnology (2019FBZ05).

---

**Fig S1.**
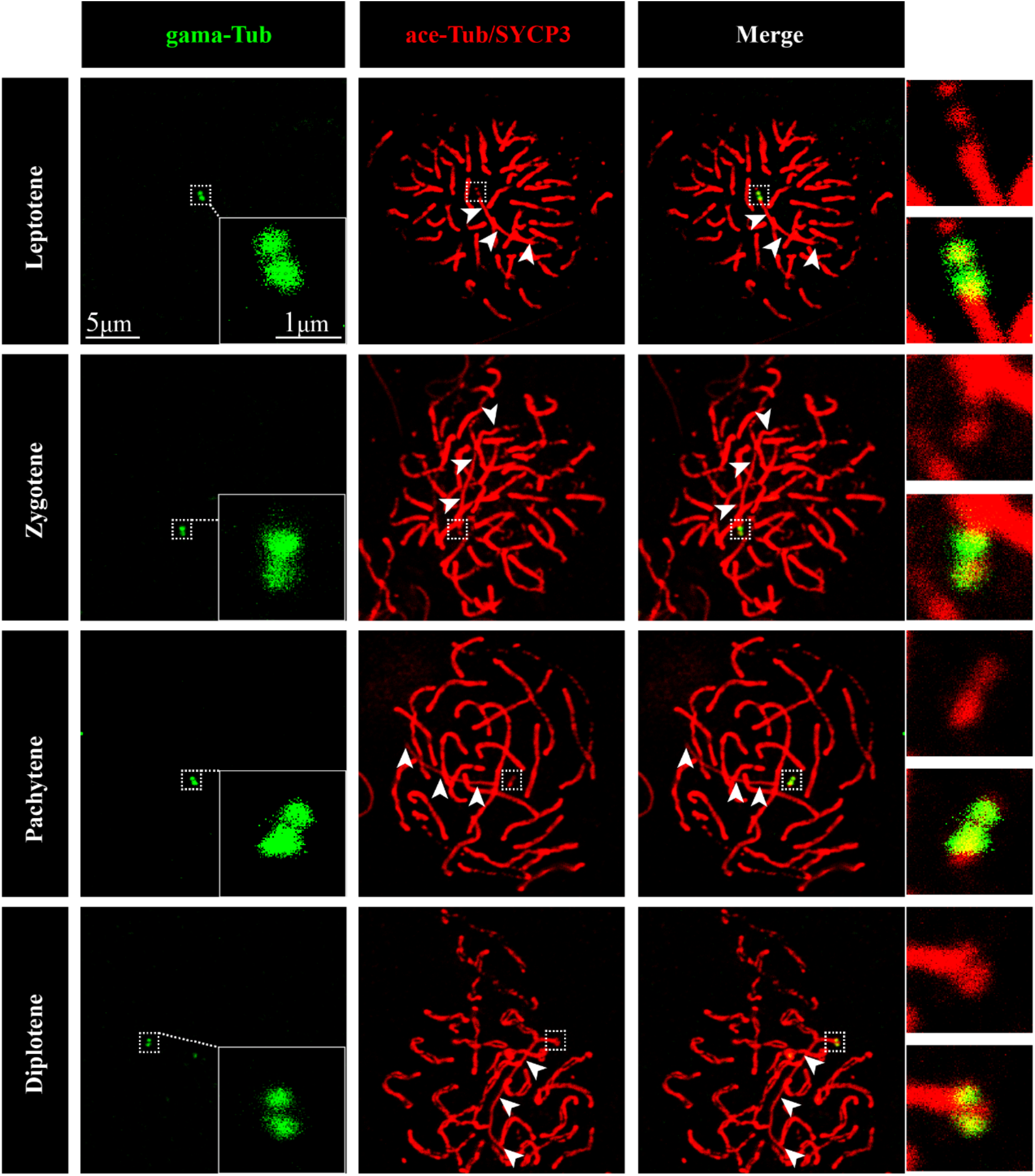
Cilia of primary spermatocyte at different stages. Confocal images showing basal bodies and synaptonemal complexes labeled with anti-γ tubulin (green) and anti-SYCP3 (red) antibodies at different stages of primary spermatocytes as indicated. Because both SYCP3 and acetylated-tubulin antibodies were raised in rabbit, spermatocyte cilia were also labeled in red (white arrowhead). The spermatocyte cilia can be distinguished by the basal colocalization with anti-γ tubulin antibodies.

**Fig S2.**
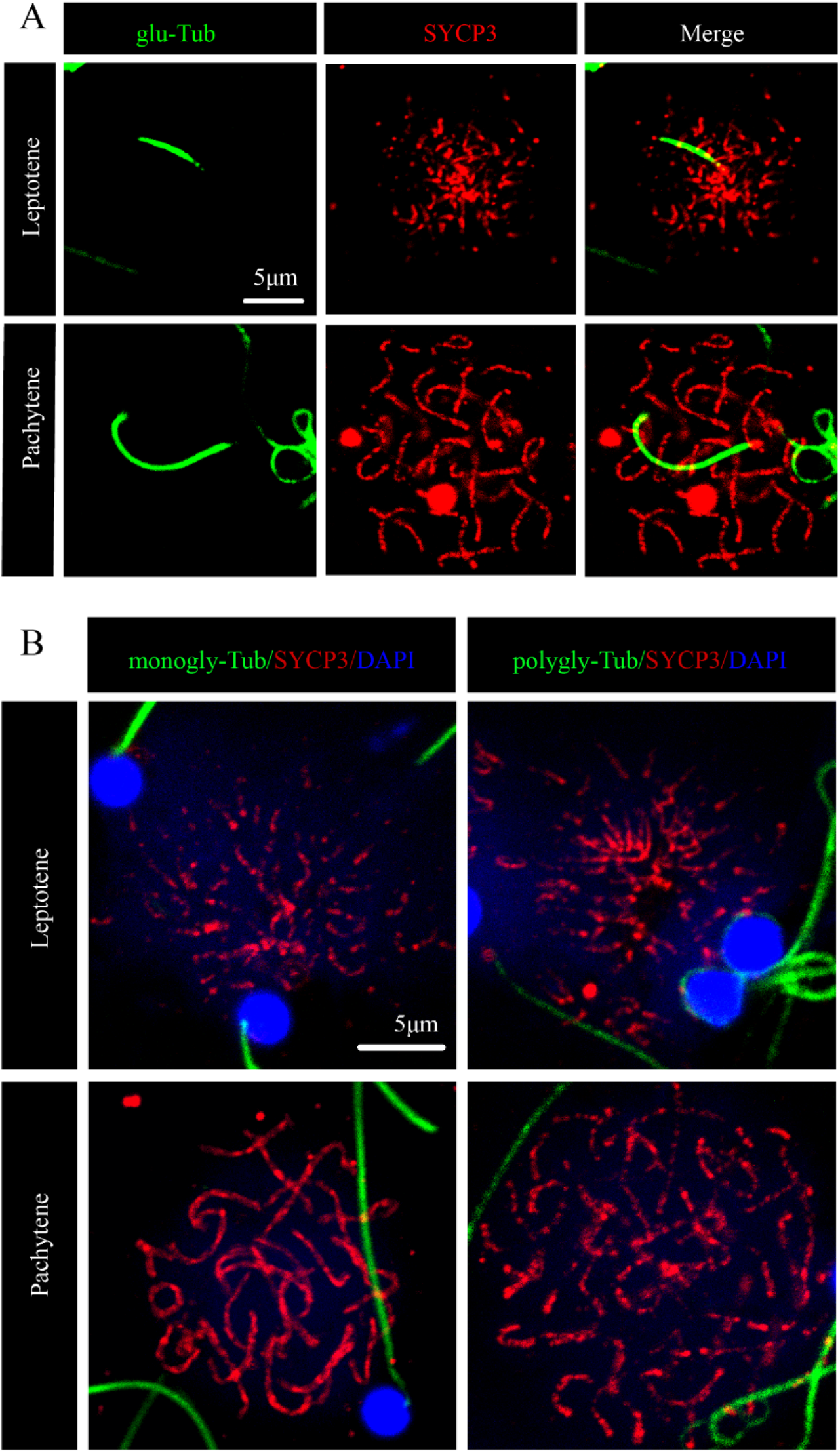
Tubulin modification inside spermatocyte cilia. (A) Confocal images showing cilia and synaptonemal complexes labeled with anti-glutamylated-tubulin (glu-Tub, green) and anti-SYCP3 (SYCP3, red) antibodies. (B) Confocal images showing staining of mono-glycylated (monogly-Tub, green) or poly-glycylated (polygly-Tub, green) Tubulin antibodies. The sperm flagella, but not spermatocyte cilia, can be labeled with these antibodies.

**Fig S3.**
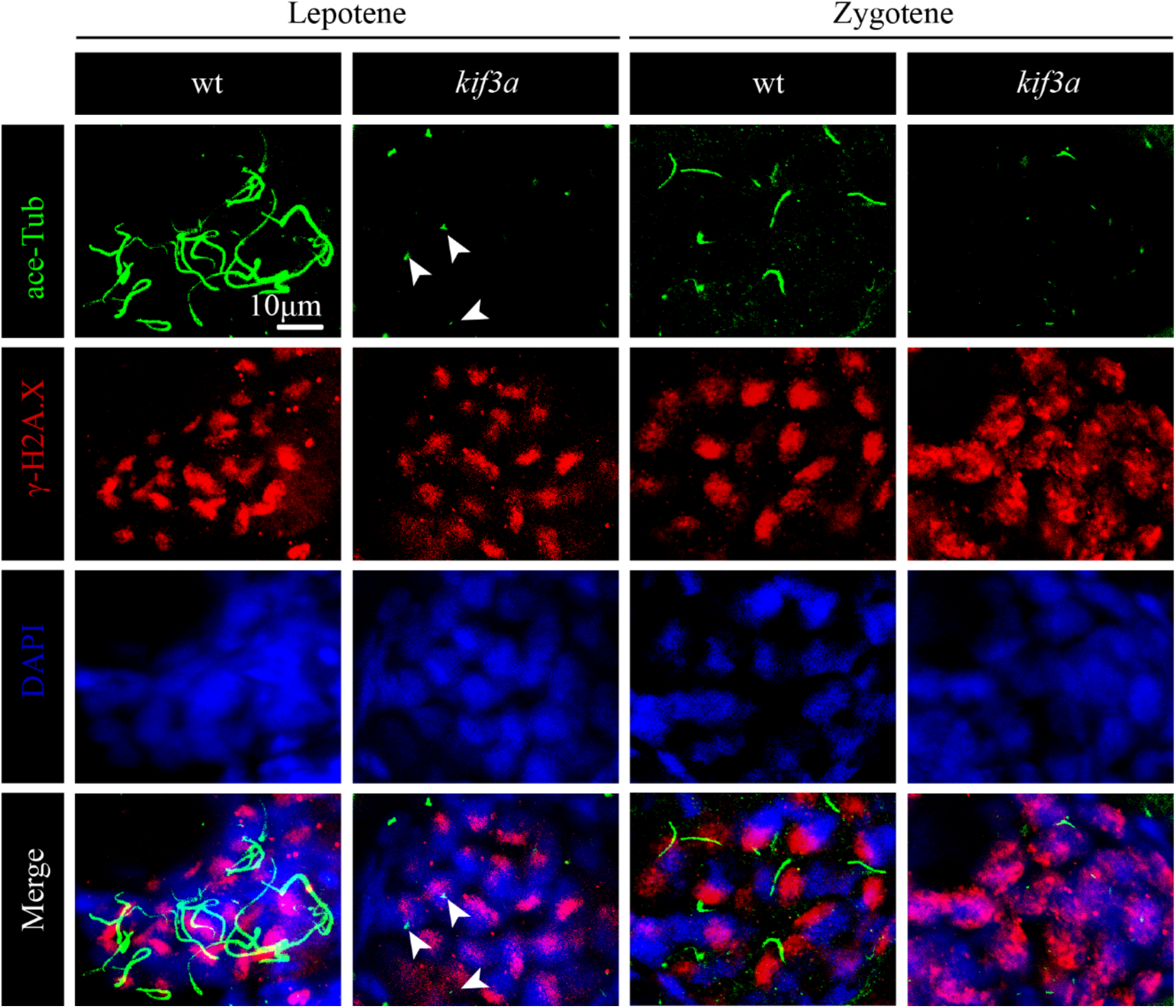
Cilia were present in primary oocytes. Staining of γ-H2A.X (red) and acetylated tubulin (green) in the oocytes of wild-type and *Tg (kop:cas9-UTRnanos3; U6:3xsgRNA-kif3a)* double transgenic fish. Arrowheads indicate basal bodies left in the mutant oocytes.

**Fig S4.**
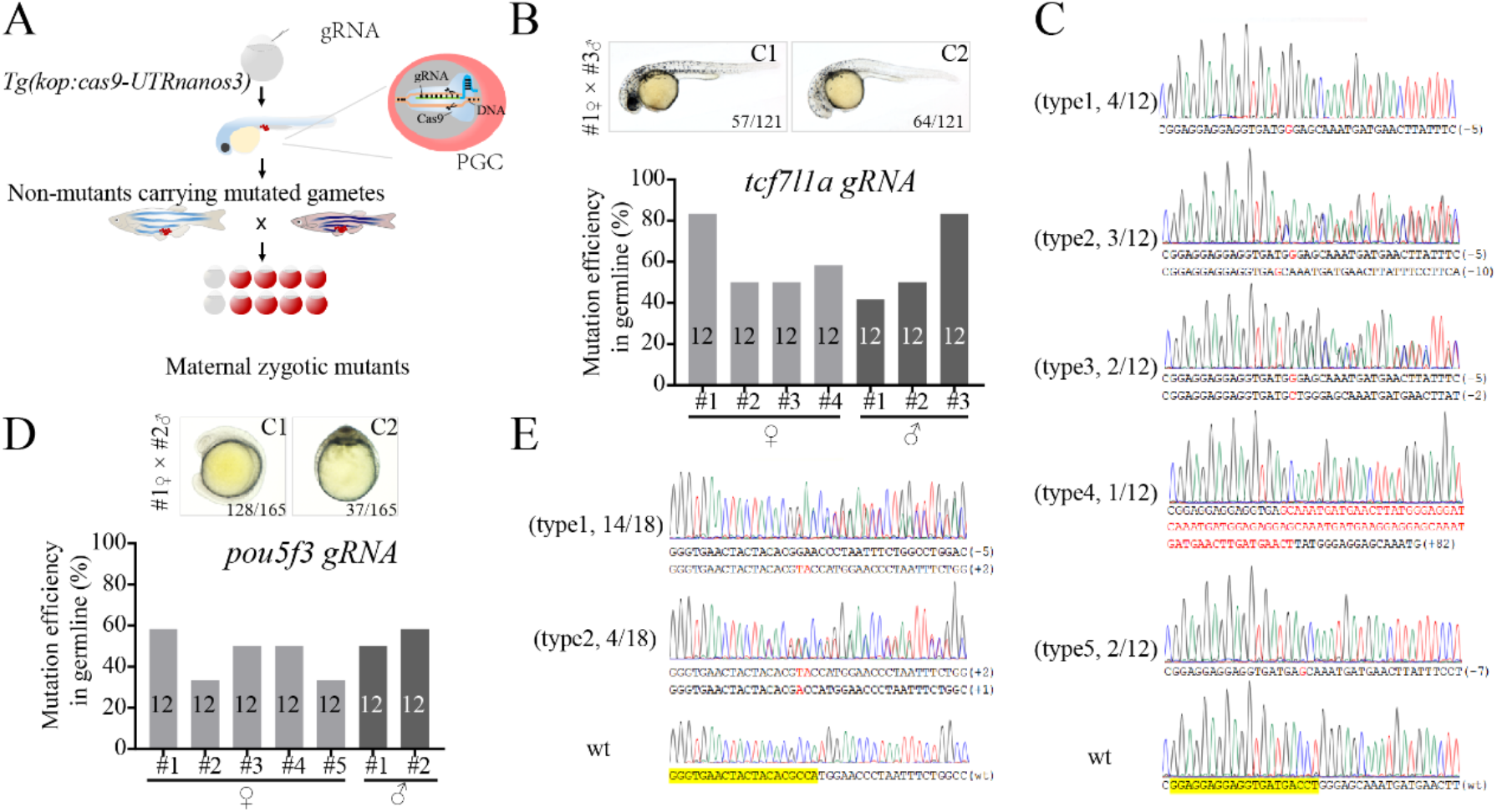
Generation of maternal-zygotic mutants via PGC-specific Cas9 transgenic line. (A) Schematic workflow showing process of generating MZ mutants using PGC-specific Cas9 expressing *Tg(kop:cas9-UTRnanos3)* embryos. (B) Mutation efficiencies of gametes and the phenotypes of offspring of *tcf7l1a* sgRNA injected *Tg(kop:cas9-UTRnanos3)* fish. C1 shows the WT like phenotype, C2 shows complete loss of eyes. The mutation efficiencies were calculated by the number of mutated heterozygotic embryos from crossing between F0 and wild type fish. A total of 12 embryos were genotyped from each cross. (C) The genotype of MZ*tcf7l1a* with C2 phenotype in panel B. (D) Mutation efficiencies of gametes and the phenotypes of offspring of *pou5f3* sgRNA injected *Tg(kop:cas9-UTRnanos3)* fish. C1 shows the WT like phenotype, C2 shows the phenotype of MZ*pou5f3*. (E)The genotype of MZ*pou5f3* with C2 phenotype in panel D.

**Fig S5.**
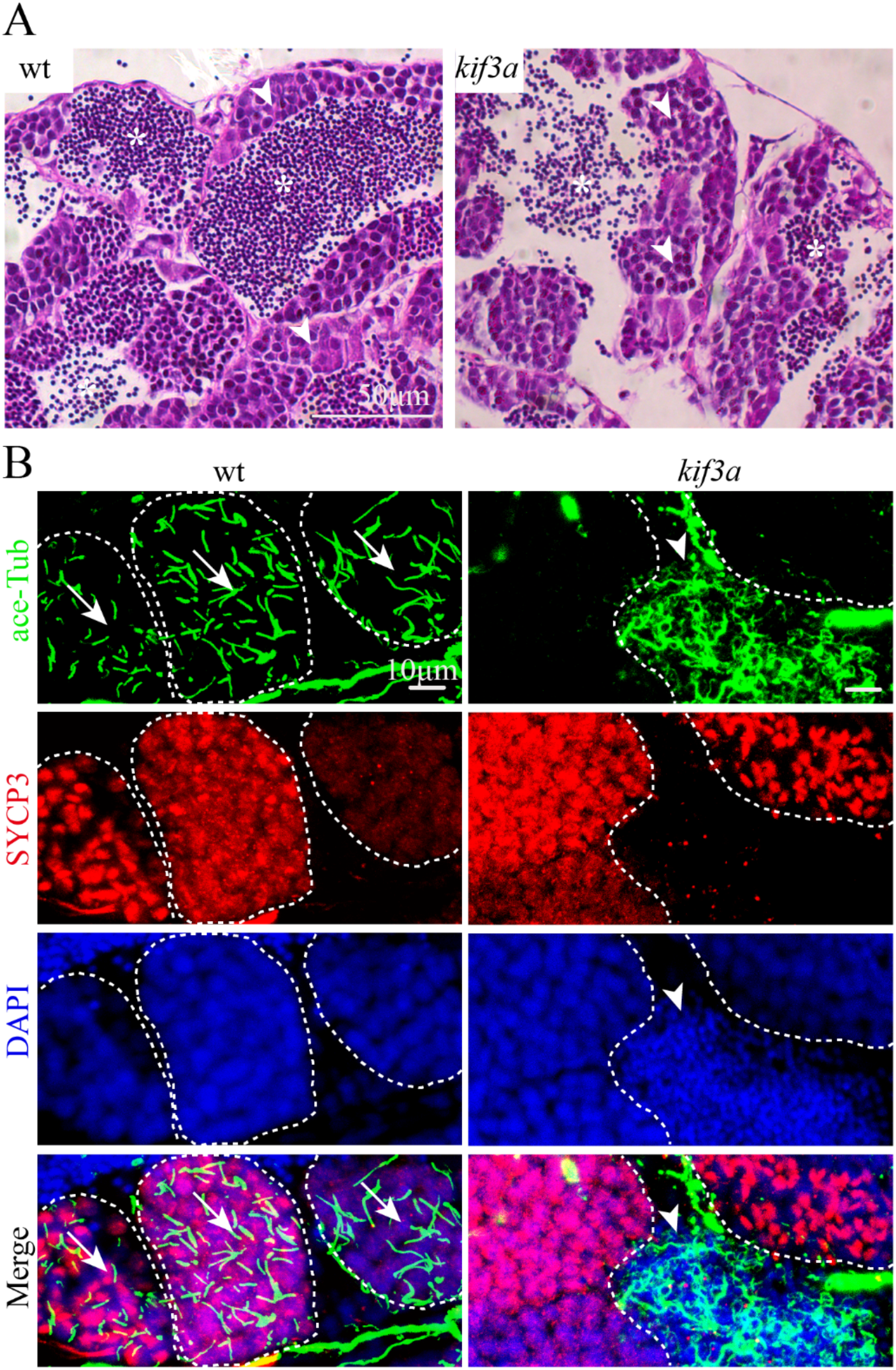
Germ cell-specific Kif3a conditional knockout inhibited the formation of spermatocyte cilia. (A) Histological analysis of testis of wild type and *Tg(kop:cas9-UTRnanos; U6:3xsgRNA-kif3a)* double transgenic fish. Arrowheads point to spermatocyte and asterisks indicate spermatozoa. (B) Staining of SYCP3 (red) and acetylated-tubulin (green) in the testis of wild-type and double transgenic fish. Arrows indicate spermatocyte cilia, while arrowheads indicate the staining of abnormal sperm flagella in the mutant testis.

**Fig S6.**
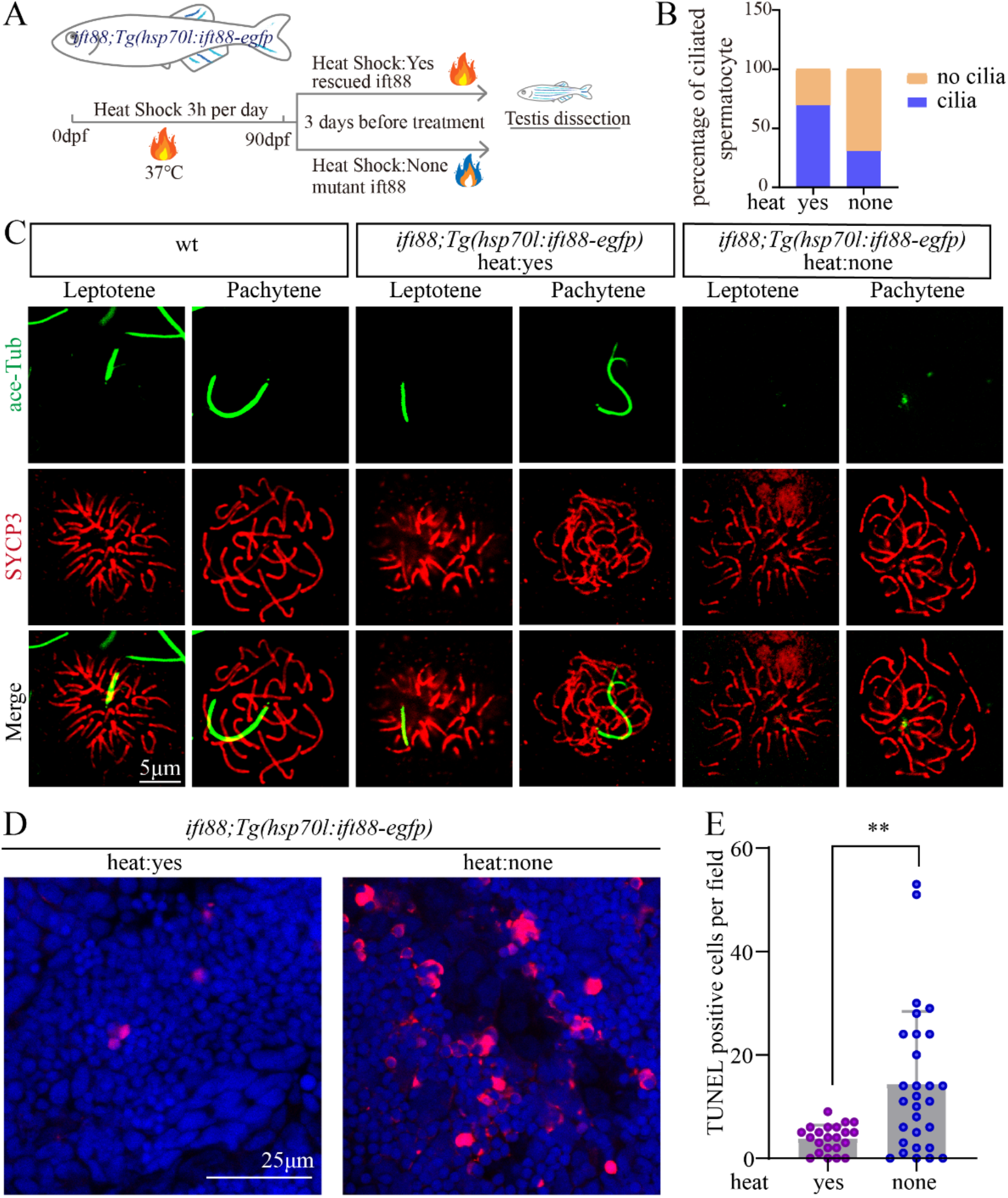
Ift88 deficiency resulted in ciliogenesis defects and apoptosis of spermatocytes. (A) Schematic workflow showing the strategy of generating *Tg(hsp70l:ift88-egfp)* transgene to rescue *ovl(ift88)* mutants. (B) Statistical analysis of the percentage of ciliated spermatocytes in heat-shocked and non-heat-shocked *ift88;Tg(hsp70l:ift88-egfp)* transgenic fish. (C) Confocal images showing cilia and synaptonemal complexes labeled with anti-acetylated tubulin (green) and anti-SYCP3 (red) antibodies on primary spermatocytes from different fish as indicated. (D-E) Confocal images showing apoptotic cells stained by TUNEL assay (D) and the statistical results (E) of TUNEL positive cells in heat-shocked and non-heat-shocked fish as indicated.

**Fig S7.**
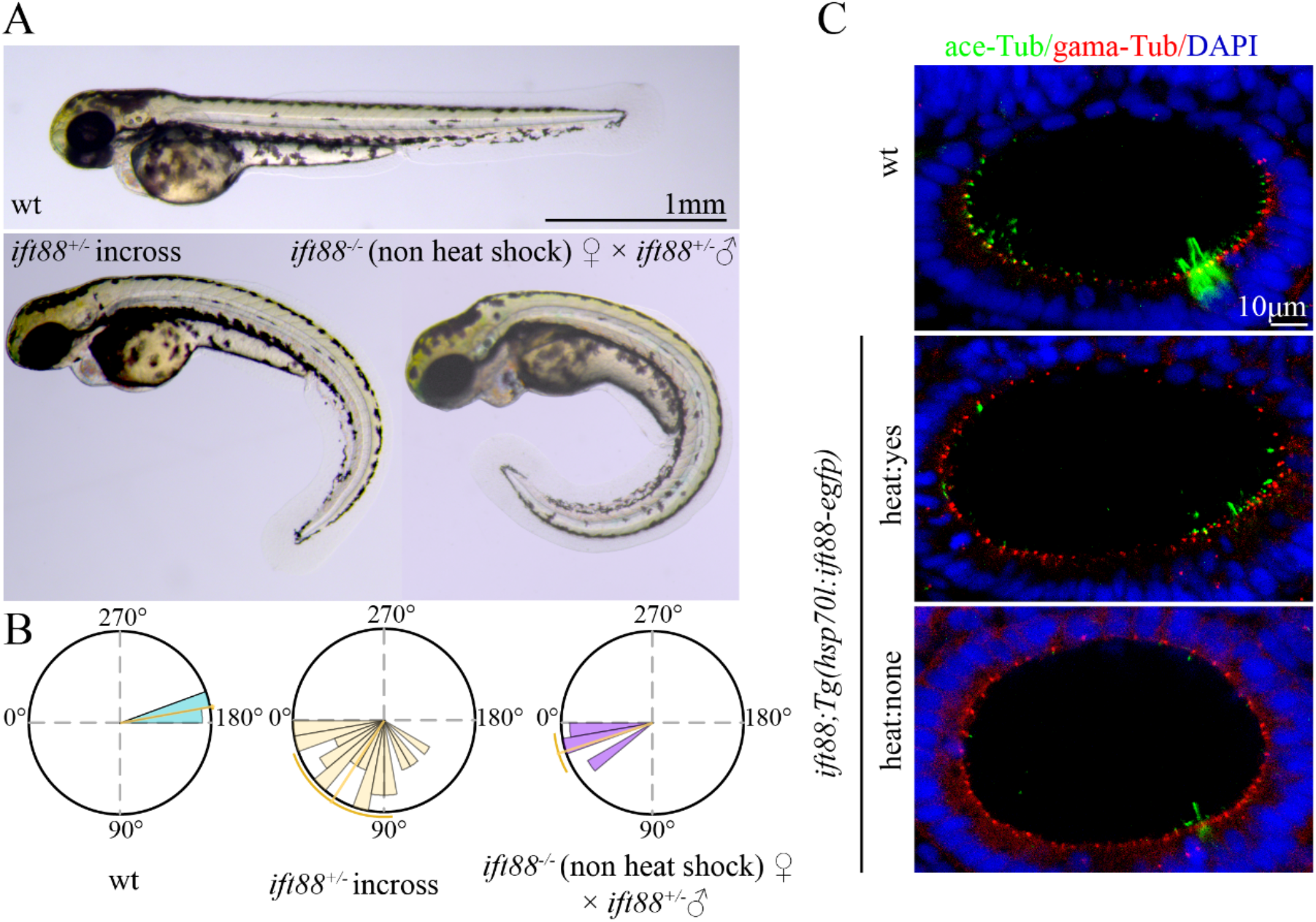
Phenotypes of *ift88* mutants. (A) External phenotypes of wild type and *ift88* mutants generated from different crosses as indicated. (B) Statistical analysis of the angles of body curvature in wild type and *ift88* mutants as indicated. (C) Confocal images showing cilia in otic vesicle of 18-somite stage wild type and *ift88* mutants as indicated.

